# Metabolic complexity increases adaptability

**DOI:** 10.1101/2023.04.17.537065

**Authors:** Claus Jonathan Fritzemeier, Sajjad Ghaffarinasab, Felix Lieder, Balázs Szappanos, Florian Jarre, Balázs Papp, Csaba Pál, Martin J. Lercher

## Abstract

Bacteria show a striking ability to adapt to new environments through horizontal gene transfer. Anecdotal evidence suggests that some bacteria are more adaptable than others: *e*.*g*., intestinal *E. coli* frequently spin off pathogenic strains adapted to other human tissues, while gastric *Helicobacter pylori* do not. However, it is unclear what determines the ability of individual strains or species to adapt to new environments. Here, we use pan-genome scale modeling to explore the ability of 102 different unicellular organisms to adapt to each of 5000+ diverse nutritional environments. While the small metabolic systems of specialized endosymbionts typically require 50+ additional metabolic reactions to adapt to new environments, different strains of the generalist *E. coli* require on average less than 5 new reactions. Thus, there is a positive feedback between metabolic complexity and adaptability, contrary to speculations that complex systems are generally less adaptable.

## Introduction

Many unicellular organisms show an astounding ability to adapt to new environments (Brooks, Turkarslan, Beer, Lo, & Baliga, 2011). Different phylogenetic lineages differ widely in the frequency with which they give rise to new strains or even new species, but it is currently unclear what determines these differences. The splitting off of new lineages will often be adaptive, with the new lineage specializing to a different life style or environment. Among bacteria, such specialization is typically accompanied by the loss of now superfluous genes from the genome and the acquisition of additional genes via horizontal gene transfer (HGT, also termed lateral gene transfer) (Koonin & Wolf, 2008; Pal, Papp, & Lercher, 2005). Examples in point are the loss of a majority of metabolic genes in the endosymbiotic bacterium *Buchnera* (Pal et al., 2006) and the acquisition of additional metabolic pathways by pathogenic *E. coli* strains to survive in the human urinary tract (Alteri, Smith, & Mobley, 2009). As a consequence of these evolutionary dynamics, bacterial pan-genomes can be partitioned into core genes (found in almost all strains), shell genes (found in several strains), and cloud genes (restricted to a single strain) (Koonin & Wolf, 2008).

Bacterial strains of the same species often differ widely in their metabolic capabilities. For example, a study on *E. coli* found that individual strains could grow in between 437 and 624 of the tested environments (Monk et al., 2013). Based on such differences, lineages can be categorized as metabolic generalists or specialists. A prolonged reduction in environmental complexity – such as experienced by a generalist bacterium becoming a permanent endosymbiont – causes a corresponding reduction in metabolic complexity, which can be predicted quantitatively from genome-scale metabolic modeling (Pal et al., 2006). That bacterial evolution appears to organize itself into short bursts of innovation followed by long phases of genome reduction (Wolf & Koonin, 2013) indicates that the inverse process – a specialist evolving into a generalist – is comparatively rare.

In previous work (Szappanos et al., 2016), we utilized metabolic simulations to show that the standard lab strain *E. coli* K-12 can adapt to most previously unviable nutritional environments by acquiring at most three additional enzymes and/or transporters via HGT. In many cases, different new environments required the acquisition of overlapping gene sets. We found that complex metabolic innovations can evolve via the successive acquisition of individual biochemical reactions, where each confers an additional benefit for the utilization of specific nutrients. This observation indicates an important role of exaptations in metabolic evolution, where stepwise metabolic niche expansion can lead to a substantial acceleration of adaptation (Szappanos et al., 2016).

However, multiple genes can also be acquired simultaneously via horizontal gene transfer. Successful transfer events of DNA in *E. coli* appear to co-transfer at most 30kb of DNA (Pang & Lercher, 2017). A reconstruction of the ancestral metabolic networks of 53 *E. coli* strains showed that all metabolic innovations identifiable *in silico* in this lineage indeed arose through the acquisition of a single DNA segment <30kb on one of the branches of the phylogeny. At the same time, around 10% of innovations relied on the exaptation of acquisitions on earlier branches of the strain phylogeny (Pang & Lercher, 2019).

These findings demonstrate that complex innovations can evolve – and have indeed evolved in *E. coli –* without the need to resort to neutral explorations of phenotype space. Such neutral explorations had been suggested earlier as an important facilitator of adaptation (Barve & Wagner, 2013), but the corresponding non-adaptive evolution is expected to be extremely slow, and no direct empirical support has been identified for this scenario in bacteria (Szappanos et al., 2016). Thus, theoretical, computational, and comparative genomics considerations indicate that bacterial evolution can be understood purely from a consideration of adaptive processes (Szappanos et al., 2016).

Compared to many other species, the generalist *E. coli* features a rather complex metabolic network. R. A. Fisher and followers believed – although without direct empirical evidence – that complexity hinders adaptation, due to increasing pleiotropic constraints (Fisher, 1930; Orr, 2005). Here, we argue the opposite. Prompted by previous evidence for a broad adaptability of the generalist *E. coli* (Pang & Lercher, 2019; Szappanos et al., 2016) and simulations of abstract representations of artificial reaction networks (Maslov, Krishna, Pang, & Sneppen, 2009), we hypothesize that bacteria with more complex metabolic systems might be more adaptable than specialists.

Below, we explore this hypothesis by studying how metabolic network size affects the adaptability of real metabolic systems. Performing metabolic simulations on a pan genome-scale, we show that the ease with which microbes adapt to new environments varies widely between species, with metabolic specialists typically requiring an order of magnitude more gene acquisitions than generalists adapting to the same environment. The increased adaptability of generalists is emphasized by their much higher potential for collateral adaptations, *i*.*e*., the ability to grow in additional, non-selected environments due to ecologically unrelated previous adaptations. Specialist species, on the other hand, have largely lost their adaptive potential. If they do adapt, however, they show a stronger tendency of exaptation, *i*.*e*., they are more likely to re-use previously acquired enzymes and transporters for later adaptations.

## Results and Discussion

### Construction of a pan-genome scale metabolic supermodel from organism-specific models

To allow coherent simulations of metabolic network expansion through HGT, we first created a pan genome-scale metabolic supermodel that contained all examined organism-specific metabolic networks as submodels. The supermodel built from 102 organismal metabolic models contains 16,018 unique reactions and 7,551 unique metabolites. **Fig. 1** shows the sizes of the organism submodels included. Most metabolites are assigned to the compartments extracellular (e), periplasm (p), and cytosol (c) (**Suppl. Fig. S2**). Several additional compartments in the supermodel originate from the contributions of four eukaryotic organisms (*Chlamydomonas* (iRC1080), *Saccharomyces cerevisiae* (iMM904, iND750), *Phaeodactylum tricornutum* (iLB1027_lipid)) and the cyanobacterium *Synechocystis* sp. PCC 6803 (iJN678). For the well-studied *Escherichia coli* str. K-12 substr. MG1655, five different models were included. We also included metabolic models for 55 other *E. coli* and *Shigella* strains (Monk et al., 2013). Further details about the organisms and metabolic models included are listed in **Suppl. Table S1**.

**Figure 1.**
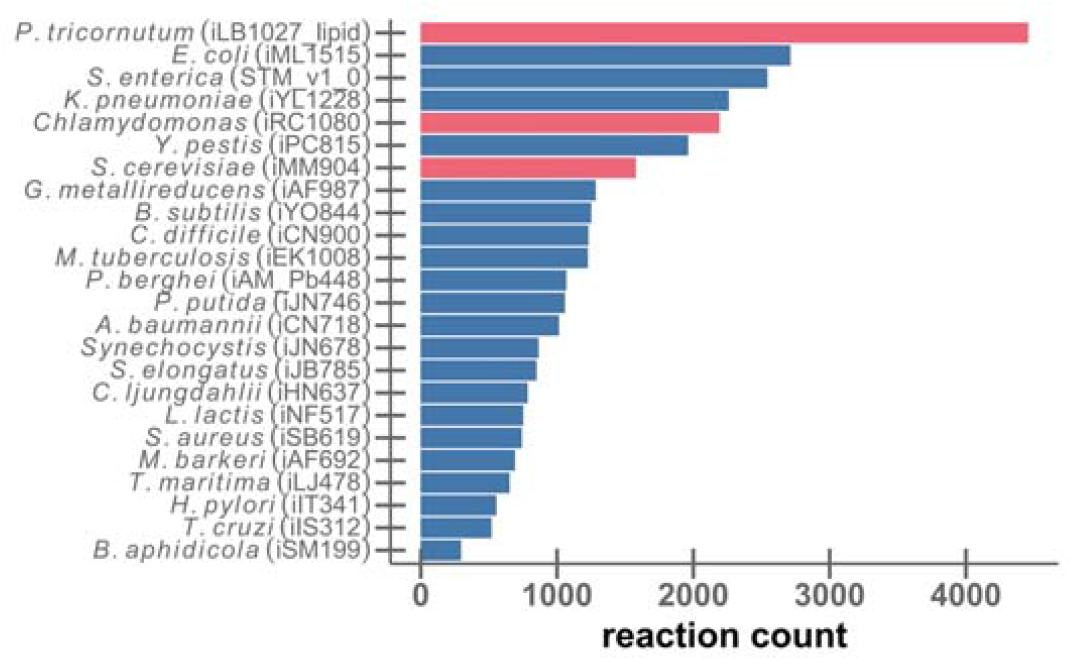
The models included represent a broad range of metabolic complexity. The bars show the number of metabolic reactions for each model that contributed to the pan-genome-scale supermodel. Only one representative strain and model is shown for each species. Red bars indicate eukaryotic models. **Suppl. Fig. S1** shows the corresponding information for additional strains and models.

Branching points in metabolic networks occur when multiple reactions produce and/or consume the same metabolite. One function of such branching points is to link alternative pathways to central metabolism. As expected, we find that larger metabolic networks tend to be less linear, i.e., they contain a lower proportion of metabolites that are consumed and produced by less than three reactions (Spearman’s ρ=-0.42, *P*=0.04; **Suppl. Fig. S4**).

We used flux balance analysis (FBA) (Orth, Thiele, & Palsson, 2010; Watson, 1984) to estimate the ability of each submodel to grow in each of a large number of nutritional environments. To make the results comparable, we used the same general biomass reaction for all organism-specific submodels, *i*.*e*., each metabolic system was required to produce the same metabolic precursors for cellular growth (Methods). We examined two sets of nutritional environments: one set that largely contains typical wet lab growth media (Henry et al., 2010), including those assayed in the Biolog phenotyping system; and another set of random minimal media, each comprising a combination of carbon, nitrogen, sulfur, and phosphorus sources plus trace elements.

As most models cannot grow in any of the random minimal environments, we checked whether all models can grow in a medium that supplies all possible nutrients. Only three models are not viable in this maximally rich condition: the hyperthermophilic bacterium *Thermotoga maritima* (iLJ478), the parasitic protozoon *Trypanosoma cruzi* Dm28c (iLS312), and the endosymbiotic bacterium *Buchnera aphidicola* (iSM199). This is because the general biomass objective function contains more amino acids than the original biomass functions of these models. We chose not to exclude these models from further analyses, as the ability of extreme specialists to adapt to new environments is one of the questions we aim to explore.

As shown in **Fig. 2**, the minimal random environments are too restricted for most modeled organisms and hence provide limited insights into the growth of the submodels in the real world. In contrast, almost all submodels can grow in at least some of the wet lab environments, with the most versatile model – *E. coli* – growing in 36% of wet lab media (**Fig. 2**). The distribution of the fraction of viable wet lab environments across submodels is bimodal (**Suppl. Fig. S5**), naturally dividing these organisms into generalists and specialists; we set the dividing line at growth in 20% of assayed media. As expected, the same three organisms unable to grow in the full medium are also unable to grow in any wet lab environment. To guard against any biases introduced by the general biomass function, we repeated this analysis with using the generation of energy (conversion of ADP to ATP) as the objective function, with qualitatively similar results (**Suppl. Fig. S6)**.

**Figure 2.**
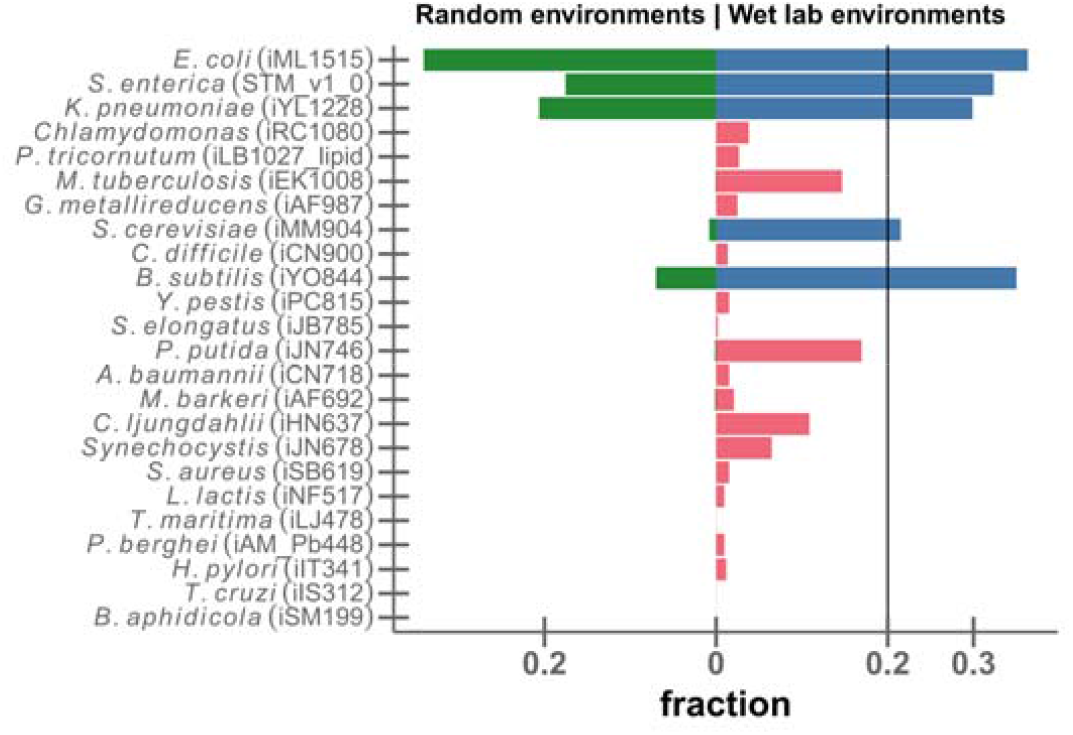
The fraction of viable environments differs widely across submodels, both for random minimal environments (green bars to the left) and for common wet lab environments (blue and red bars to the right). The vertical line indicates the mean fraction of viable wet lab environments; we use it as the threshold for partitioning metabolic systems into generalists (blue) and specialists (red). Models are ordered top to bottom by decreasing genome size. **Suppl. Fig S6** shows the corresponding results when using energy production instead of biomass production as the objective function.

### More complex networks are more adaptable

We next quantify the difficulty for an organism to adapt to new environments. For each submodel and each environment in which it is currently unable to grow, we identified the minimal number of reactions that have to be added to produce biomass; below, we refer to this number as the *added reactions*. The distribution of added reactions per wet lab environment varies widely across organisms (**Fig. 3a**, including only one representative for *E. coli*). Results are quantitatively similar when considering random instead of wet lab environments (**Suppl. Fig. S7**), and qualitatively similar when using energy generation instead of biomass production as the objective function (**Suppl. Fig. S8**).

**Figure 3.**
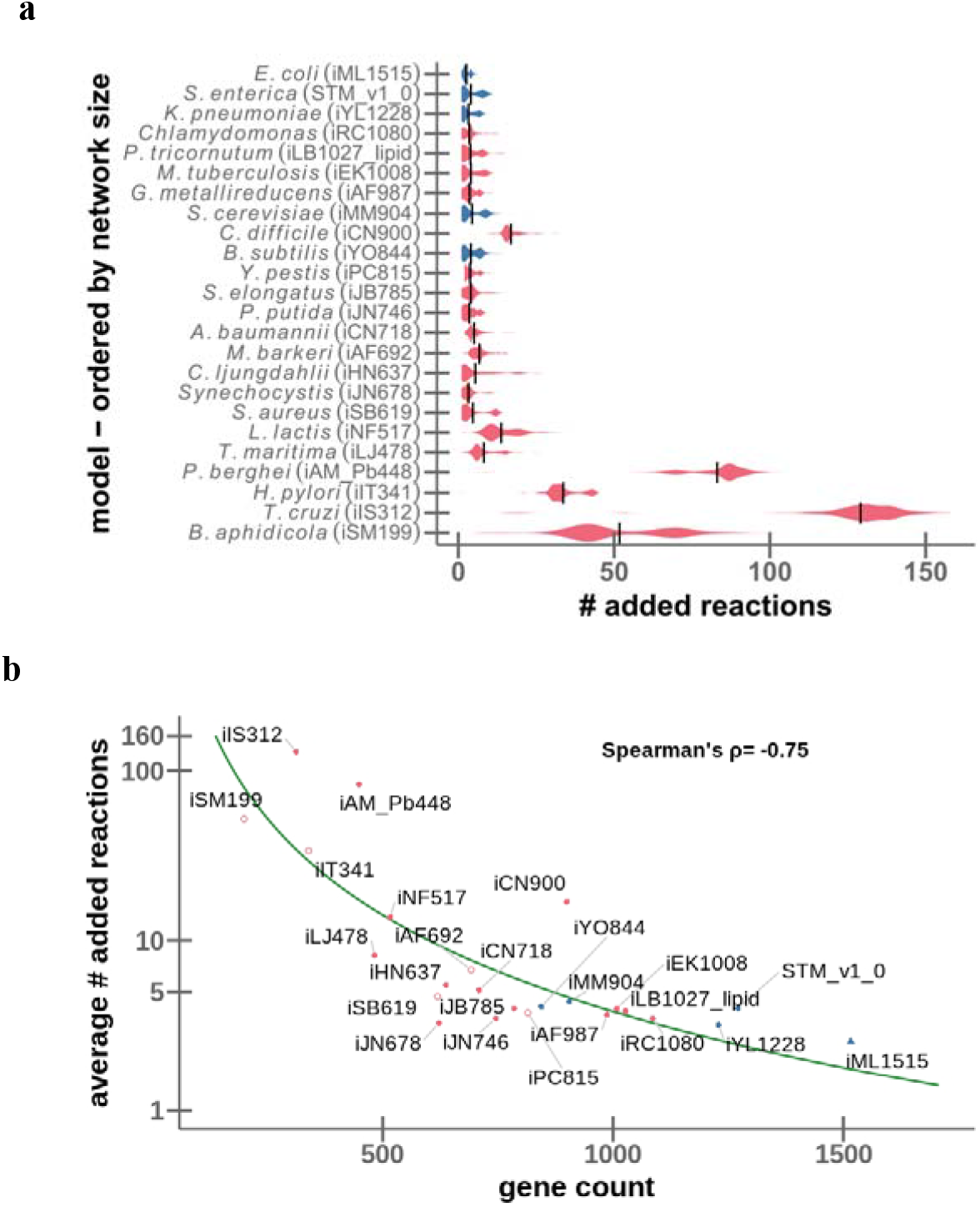
The number of additional reactions required for adaptation decreases with increasing genome size. **(a)** Distributions of added reactions, summarized as violin plots. The height at each point in a “violin” indicates the local density of the distribution for the given model. Models are ordered top-down by decreasing size. **(b)** The average number of added reactions (log scale) plotted against metabolic gene count for each model. The green line shows the best fitting power law, *added reactions* = *a* × (*gene count*)^*b*^. In both panels, colors distinguish specialists (red) and generalists (blue). Organisms with known auxotrophies are shown as open circles. The 62 *E. coli* strains are represented by the iML1515 model (blue triangle) only. For this figure, only wet lab environments are considered.

The four smallest and most specialized metabolic networks require the largest number of added reactions to adapt to new environments. The endosymbiont *Buchnera aphidicola* needs to add on average 51.7 reactions to reach new environments. Similarly, the pathogen *Helicobacter pylori*, which exclusively lives in human stomachs, needs on average 33.7 additional reactions. *Plasmodium berghei*, which is a protozoan parasite that causes malaria in rodents, requires on average 83.0 reactions to be viable in a new environment. Finally, the parasite *Trypanosoma cruzi Dm28c* requires on average 129.1 reactions. All four organisms are highly specialized to one or a few specific, stable environments. Accordingly, their metabolisms show very little flexibility, reflected in very small numbers of metabolic genes (*B. aphidicola:* 199 metabolic genes out of a total of 517 genes (Shigenobu, Watanabe, Hattori, Sakaki, & Ishikawa, 2000); *H. pylori:* 341 metabolic genes out of 1590 total genes (Tomb et al., 1997); *T. cruzi*: 312 metabolic genes out of 1430 (De Pablos & Osuna, 2012); *P. berghei*: 448 metabolic genes out of 5216 (Otto et al., 2014)). At the other end of the spectrum in **Fig. 3a** is *E. coli*: the standard lab strain K12 (iML1515) requires on average 2.55 and at most 6 reactions to adapt to any of the tested environments.

Although it is likely that many properties influence the ability of a metabolic system to adapt to new nutritional environments, network size alone explains 56% of the variance across all assayed models (**Fig. 3b**; Spearman’s ρ = −0.75, *P* = 4.1×10^−5^). The solid line in **Fig. 3b** shows the best-fitting power law, *added reactions* = *a* × (*gene count*)^*b*^. The best-fitting exponent is *b* = 2.87 (95% CI = [3.47, 2.28]), which is slightly larger than the *b* = 2 expected from abstract models of metabolic network expansion (Maslov et al., 2009).

The dataset contains *E. Coli* models of various sizes, with between 904 and 1,516 metabolic genes. These mostly differ only marginally in their adaptability (**Suppl. Fig. S9**): the average number of added reactions for generalist *E. coli* (including nine strains with auxotrophies) lies between 2.40 and 2.97, while the average number of added reactions for specialist *E. coli* ranges from 2.30 to 5.22. The outlier requiring the largest number of additional reactions is *E. coli* DH1 (iEcDH1_1363; **Suppl. Fig. S9**), which is auxotrophic for thiamine (Meselson & Yuan, 1968) due to the loss of a complete operon (Monk et al., 2013). Similar to the picture across species (**Fig. 3b**), and despite the low variation in adaptability, we find a substantial negative correlation between the average number of added reactions and gene count when comparing different *E. coli* strains (Spearman’s ρ=-0.60, *P*=6.3×10^−6^, excluding strains with auxotrophies).

### Specialists often re-use gained reactions in later adaptations

If an organism adapts to a given environment by acquiring a matching set of metabolic reactions, it can happen that the same reactions now also facilitate growth in another environment, where the organism was unviable before. With few exceptions, such collateral, unselected-for adaptation happens more frequently for generalists than for specialists across species; this trend is reversed when comparing different *E. coli* strains, possibly because the repair of auxotrophies facilitates growth in multiple environments (**Suppl. Fig. S10**; see also Refs. (Barve & Wagner, 2013; Hosseini & Wagner, 2016)).

But even if the reactions acquired to adapt to environment *A* do not provide immediate access to environment *B*, they may still provide a subset of the reactions required to adapt later to this second environment. We quantify the propensity to profit from adaptations in this way with an *exaptation index* (see Methods). One might hypothesize that while specialists show little collateral adaptation, they may show a high potential for step-wise exaptation, for example if added reactions remove an auxotrophy. As expected from this hypothesis, **Suppl. Fig. S11** shows that we indeed tend to find higher exaptation indices for specialists than for generalists; moreover, the propensity for such exaptations is higher for specialists with small genomes compared to specialists with larger genomes.

## Conclusions

Adaptations arise by extensions of existing phenotypes and genotypes. In specialists with small genomes, adaptation to new ecological niches is typically difficult, as it demands the simultaneous acquisition of multiple mutations/genes. As a consequence, specialists with simple genomes may often be evolutionary dead-ends. The smallest and most specialized metabolic systems, those of *Buchnera aphidicola* (an endosymbiont of aphids), *Trypanosoma cruzi Dm28c* (an internal human pathogen), and *Helicobacter pylori* (an endosymbiont of the human stomach), are trapped in their endosymbiotic life style, having all but lost their adaptive potential. The opposite is true for organisms with complex genomes – such as *E. coli* – whose larger “toolboxes” (Maslov et al., 2009) can more easily be extended for novel tasks.

The observed relationship between metabolic network size and adaptability leads to a positive feedback between complexity and evolvability. This conclusion is the exact opposite of what is suggested by Fisher’s geometric model (Fisher, 1930; Orr, 2005). Fisher’s model supports the idea that more complex systems are less likely to adapt through natural selection. In support of this idea, it has been observed that genes encoding proteins involved in many protein-protein interactions are less likely to be horizontally transferred than genes encoding less highly-connected proteins (Cohen, Gophna, & Pupko, 2011; Jain, Rivera, & Lake, 1999). This effect might be expected, as the interaction between two proteins requires the co-evolution of the amino acid sequences directly involved in the contact, and hence a protein encoded by a newly acquired gene may not bind sufficiently strongly to existing proteins of the host. Conversely, different enzymes that interact in a metabolic network perform their molecular functions independently, and their amino acid sequences do not need to be finetuned with respect to each other. This line of argument suggests that metabolic genes with high connectivity may be integrated easily into an existing network, while genes with high connectivity in the protein-protein interaction network may not.

However, while the amino acid sequences of different enzymes may be independent of each other, their expression has to be coordinated precisely. Thus, finetuning is necessary also for the integration of metabolic genes into an existing network, although the adjustments must occur in terms of regulatory changes rather than amino acid sequence changes. That metabolic complexity and protein-protein interaction complexity appear to have opposite effects on adaptability might then be explained by faster adaptive evolution of gene expression compared to amino acid sequences (Lenski, 2017; Lozada-Chavez, Janga, & Collado-Vides, 2006).

Exaptation – the utilization of metabolic genes acquired in previous adaptations for adaptive purposes in a new environment – plays an important role in the adaptation of both generalists and specialists, although in different ways. Generalist species, but not specialist species, show a high degree of collateral adaptation, *i*.*e*., previous adaptations often enable growth in environments other than those experienced by the organisms’ ancestors (Barve & Wagner, 2013). Conversely, specialist species that acquire new metabolic genes in the adaptation to one environment are more likely to re-use (exapt) these genes in later adaptations to other environments; thus, stepwise metabolic niche expansion will play an even stronger role in the adaptation of specialists than previously observed for the generalist *E. coli* (Szappanos et al., 2016), and might thus be the facilitator of rare genome expansions (Koonin & Wolf, 2008).

## Materials and Methods

### Supermodel generation

We started with 109 genome scale models (GSMs) downloaded from the BiGG database (Schellenberger, Park, Conrad, & Palsson, 2010). We removed seven models of multicellular eukaryotes. As we are specifically interested in variations in metabolic model size and as the BiGG database contains only few species with very small metabolic systems, we added the model for *Buchnera aphidicola* str. APS (Macdonald, Lin, Russell, Thomas, & Douglas, 2012). Thus, 102 GSMs (termed “submodels” in this work) contributed to the supermodel (**Suppl. Table S1**). As a preprocessing step, we checked whether reactions and metabolites from different submodels but with the same IDs represented the same biochemical reaction; if not, we renamed them. Reactions were compared on the basis of stoichiometry and reversibility, while metabolites were compared on the basis of their chemical formulas if these were available.

A preliminary supermodel was formed as the union of the reactions and metabolites from all submodels. As detailed below, we then curated this preliminary model by ensuring mass balance and by removing energy generating cycles (EGCs) (Fritzemeier, Hartleb, Szappanos, Papp, & Lercher, 2017). While each individual model passes these quality checks, the reactions in the merged supermodel may be combined in ways that violate thermodynamic laws or the mass balance. Mass balance is considered first, because proper mass balance is a requirement for the EGC removal. The final supermodel was thus mass-balanced and had no energy-generating cycles.

### Correction of mass balance

Mass balance of a reaction is generally ensured by contrasting all atoms of the educts and all atoms of the products. However, due to incomplete data, the mass balance for many reactions is not known; removing all reactions with uncertain mass balance would render most of the models non-functional. To circumvent this problem, only the mass balance of the exchange reactions was considered: the number of atoms of the same kind (*e*.*g*., carbon) entering the model has to equal the number of corresponding atoms leaving the model. The only reactions that allow exchange of molecules with the model environment are exchange reactions and biomass reactions. At the same time, these are the only reactions in a network that are allowed to be imbalanced. We first removed exchange reactions and biomass objective functions that contain a metabolite of unknown composition from the model, as for these we cannot guarantee mass balance. To identify potentially imbalanced reactions, we fixed the net exchange of atoms to zero. We then removed all reactions that are blocked in this situation.

### Removing erroneous energy-generating cycles

Another problem occurring when combining multiple GSMs is the formation of erroneous energy generating cycles (EGCs) (Fritzemeier et al., 2017; Szappanos et al., 2016). In GSMs, such thermodynamically impossible cycles can produce energy equivalents (*e*.*g*., by synthesizing ATP) in infinite amounts without the consumption of nutrients (Fritzemeier et al., 2017). Thermodynamics are strongly influenced by metabolite concentrations. However, GSMs consider thermodynamics only approximately through the directionality of reactions. Thus, combining two networks can cause the formation of EGCs even if the individual networks are EGC-free.

Based on a previously published algorithm (Fritzemeier et al., 2017), we constructed a greedy approach to build organism-specific EGC-free supermodels. We chose not to build one supermodel for all analyses, as the order of adding metabolic networks to the growing supermodel can affect the final model, and as we wanted to study the adaptability of each organism starting from a model from which none (or only a few) reactions had been removed.

From the preliminary, mass-balanced supermodel, we first considered the set of reactions of the focal organism and removed any EGCs present. We then iteratively added the remaining submodels, each time removing all EGCs before proceeding to the next one. The order of adding organisms was determined by the initial number of EGCs; models with fewer EGCs were always added first.

To remove EGCs, we first determined the smallest set of reactions capable of producing energy equivalents in the model. This problem was solved in previous work with the ARM MILP algorithm, but here we instead used the ARM LP algorithm (see Methods, “Active reaction minimization”). We randomly chose one reaction in this cycle; we deleted the reaction if it was irreversible, and constrained it to be irreversible in the opposite direction if it was reversible. We repeated this process until no more EGCs were present. This procedure resulted in one mass-balanced, EGC-free supermodel for each organism-specific model in our dataset.

### Active reaction minimization

Mixed integer linear programs (MILP) are frequently used to extend FBA, *e*.*g*., in ROOM (Satish Kumar, Dasika, & Maranas, 2007), gapfind, and gapfill (Shlomi, Berkman, & Ruppin, 2005). In many of these problems, the objective is active reaction minimization (ARM). The pan-genome-scale model in this work is much bigger than any genome-scale models. Current methods of minimizing the number of active reactions under flux balance constraints cannot be applied due to the exponential complexity of this problem. We here use an approximate method that leads to major speedups and minor inaccuracies. A corresponding linear approximation has also been used in combination with the Gapfill algorithm (Thiele, Vlassis, & Fleming, 2014).

We relax the following ARM MILP problem into a sequence of ARM LP^k^ *k* ∈ :{1, … *n*} for problems. We use the property of the simplex algorithm to find sparse solution vectors.

ARM MILP:

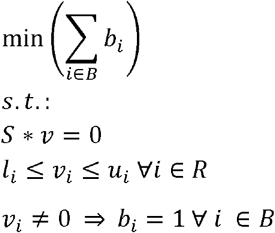

ARM LP^k^:

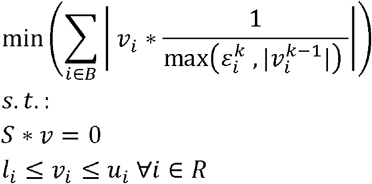

**Table 1.** Definition of variables of the ARM LP.

|  |  |
| --- | --- |
| $S \in \mathbb{R}^{\vert M\vert } \times \mathbb{R}^{\vert R\vert }$ | stoichiometric matrix |
| $v \in \mathbb{R}^{\vert R\vert }$ | Vector of fluxes |
| $l \in \mathbb{R}^{\vert R\vert }$ | Vector of lower bounds |
| $u \in \mathbb{R}^{\vert R\vert }$ | Vector of upper bounds |
| $b \in \{0,1\}^{\vert B\vert }$ | Vector of binary variables |
| $R \in \{1, \dots, m\}$ | Set of $m$ reaction indices |
| $B \subseteq R$ | Set of indices that are objective of the optimization |
| $v_i^k$ | Flux of the $i$-th reaction in the $k$-th optimization (0 if undefined) |
| $k$ | Optimization step counter |
| $n$ | Total number of optimization steps |
| $\varepsilon_i^k$ | $i$-th upper bound of weight factor in optimization round $k$ |

In this sequence of linear problems, the optimization function of the *(k+1)*-th problem is reweighted with the solution of the *k*-th problem. The initial values for 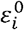are either set to one or to some positive random values. For the (*k*+1)-th optimization, we recalculated 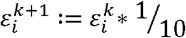.

In order to show the practical application of our linear approximation of active reaction minimization, we show the comparison between the MILP result and the LP approximation. To limit the computation time to a reasonable span, we allowed the solver for the MILP eight parallel threads per problem and a maximum time of two minutes per problem. Thus, some results are suboptimal, but the gap value accounts for the maximal possible difference to the optimal value. For the ARM LP calculations, the linear problem was solved twelve times and the best solution was kept. After every fourth optimization, *ε* was reinitialized with random values and *ν* is set as undefined.

**Suppl. Fig. S12a** shows both results in direct comparison for a total of 2830 problems we solved with the *E. coli* model iAF1260 and the standard biomass reaction. The ARM LP performs better for some problems with the non-optimal MILP solutions, *i*.*e*., with a gap greater zero. This is also the case for the exact solutions. We suspect the solver to have some numerical issues and thus to give a non-optimal solution in four MILP cases. All results were successfully verified with FBA. The differences between the pairwise results are shown in **Suppl. Fig. S12b**. For over 50% (1587 of 2830) of the problems, the ARM LP found a better or equally good optimal value. For 75% of the problems, the ARM LP solution differed by at most two reactions from the MILP ARM solution. Without a pre-specified time limit, the MILP ARM computation times vary widely (from seconds to hours), while the ARM LP problems are always solved in split seconds.

### Environmental Distances

The environmental distance is the minimal number of reactions an organism has to obtain in order to survive in an environment that did not support growth beforehand. The environment is defined as the set of nutrients available for growth, and viability is defined as the ability to produce biomass at a rate above 0.01 mmol gDW^-1^ h^-1^ This calculation depends on two major factors: the definition of the environments, *i*.*e*., the growth media, and the choice of the biomass objective function for a model.

### Sets of environments

All molecule types that can be taken up by the supermodel are potential nutrients. Environments differ by which of these potential nutrients are present. We analyzed two sets of environments. The first set (“wet lab media”) is taken from the Seed database (Henry et al., 2010) and represents wet lab growth media. The environments in the second set (“random media”) are derived from a minimal growth medium for the *E. coli* model iAF1260. Each of these environments consists of one carbon, one nitrogen, one sulfur, and one phosphorous (CNPS) source, accompanied by trace elements essential for growth (Szappanos et al., 2016). The complete set is generated by randomly choosing 5000 such combinations.

### Biomass objective functions

Each model includes the definition of at least one biomass reaction. For the further analyses, we selected one of these. The biomass reactions of 16 models were blocked in the mass balance step. These models did not have a valid biomass reaction anymore and were excluded from the analyses that were based on the model-specific biomass functions. To make the submodels comparable, we defined a general biomass reaction (based on the iAF1260 biomass reaction) that contains only a set of core metabolites shared by all organisms (ribose nucleotides; deoxyribose nucleotides; amino acids; water) and an energy dissipation term (converting ATP to ADP + Phosphate + H^+^). In addition, we also considered a biomass reaction that consists only of the energy dissipation term, thus indicating if a model is able to produce energy from the nutrients. **Suppl. Fig. S13** shows the percentages of environments (wet lab or random) in which the individual submodels can produce a non-zero flux through the different biomass reactions.

### Calculation of environmental distance with ARM LP

The mass-balanced and EGC-free, organism-specific model is formally a submodel of the organism-specific supermodel (see above). For a given environment and both the submodel and the supermodel, we used standard FBA to test if biomass can be produced above the threshold of 0.01 mmol gDW^-1^ h^-1^ (“growth”). Environmental distances were calculated for environments that support growth of the supermodel but not of the submodel. For each such environment, we used ARM LP to estimate the minimal number of reactions that have to be added from the supermodel to the submodel to facilitate growth. This procedure was performed for each combination of organism-specific model, environment, and biomass reaction.

### Collateral Adaptation Index

We defined a collateral adaptation index to quantify the probability that adaptation to one environment would lead to a “collateral” adaptation to other, unselected-for environments. For each submodel, we first identified the *n* random environments in which it cannot produce biomass (unviable environments). For each of these environments in turn, we identified the smallest set of reactions from the supermodel that have to be added to enable biomass production; these reactions define the environmental distance. We then determined in how many of the *n*-1 remaining previously unviable environments this extended model can grow. If we denote this number *m*, then the collateral adaptation index is defined as the corresponding fraction, *m* / (*n*-1). Thus, an index of 1 indicates collateral adaptation to all environments, while an index of 0 indicates no collateral adaptation. This definition was similarly described elsewhere (Barve & Wagner, 2013). To make sure that the adaptations considered are indeed collateral and not selected in the initial environment, we considered only environments that had no overlap with the source environment, *i*.*e*. none of the carbon, nitrogen, sulfur or phosphate source from the adapted environment was contained in the tested environments.

### Exaptation Index

Assume that to adapt to grow in a new environment *m*_1_, an organism needs the additional reaction set *r*_1_. To grow in a second distinct environment *m*_2_, the same organism may need the reaction set *r*_2_. The fraction of preadapted reactions can be defined as 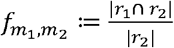. We define the exaptation index as

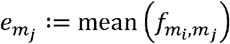

Where the mean is calculated across all *m*_i_, ∈ *M*, and *M* is the set of environments distinct from *m*_j_ in which the un-adapted submodel was unviable. For example, an exaptation index 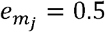 means that on average, the organism already acquired half of the reactions needed to adapt to further environments, while an exaptation index of 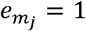 indicates collateral adaptation.

### Hardware, Software

All calculations were computed with the constraint-based modelling package “sybil” in GNU R, using IBM ILOG CPLEX as the solver. Calculations were done on a compute cluster with a peak usage of about 600 CPUs. The whole process is implemented as a pipeline reducing human interaction to a minimum. Frequent control points ensure data integrity and correctness of calculations. The code used in our simulations, as well as the corresponding results, are available on a GitLab repository at the following link: https://gitlab.cs.uni-duesseldorf.de/ghaffarinasabsharabiani/supermodel..

## Acknowledgements

We are grateful for computational support through the Zentrum für Informations-und Medientechnologie (ZIM) at Heinrich Heine University Düsseldorf. We gratefully acknowledge financial support by the German Research Foundation (DFG grants IRTG 1515 supporting CJF; CRC 680, CRC 1310 and, under Germany’s Excellence Strategy, through grant EXC 2048/1 (Project ID:390686111) to MJL), as well as through an ENORM graduate school fellowship to CJF.

## Notes

### Competing Interest Statement

The authors have declared no competing interest.

