## Supplemental Figures & Tables for "Metabolic complexity increases adaptability"

Table S1**. Organism specific models and their properties.** The models in this table are sorted in ascending order by the number of metabolic genes. The column "62 *E.coli*" indicates, whether models from the publication Monk et al. (2013).

| Model ID | Organism | Gene count | Metabolite count | Reaction count | 62 *E.coli* | Taxonomy ID | PubMed ID |
|---|---|---|---|---|---|---|---|
| e_coli_core | *Escherichia coli str. K-12 substr. MG1655* | 137 | 72 | 95 | | 511145 | 26443778 |
| iSM199 | *Buchnera aphidicola str. APS* | 199 | 298 | 297 | | | 22513857 |
| iIS312 | *Trypanosoma cruzi Dm28c* | 312 | 606 | 519 | | 1416333 | |
| iIS312_Epimastigote | *Trypanosoma cruzi Dm28c* | 312 | 606 | 519 | | 1416333 | |
| iIS312_Amastigote | *Trypanosoma cruzi Dm28c* | 312 | 606 | 519 | | 1416333 | |
| iIS312_Trypomastigote | *Trypanosoma cruzi Dm28c* | 312 | 606 | 519 | | 1416333 | |
| iIT341 | *Helicobacter pylori 26695* | 339 | 485 | 554 | | 85962 | 16077130 |
| iAM_Pb448 | *Plasmodium berghei* | 448 | 903 | 1067 | | 5821 | 29300748 |
| iAM_Pc455 | *Plasmodium cynomolgi strain B* | 455 | 907 | 1074 | | 1120755 | 29300748 |
| iAM_Pk459 | *Plasmodium knowlesi strain H* | 459 | 909 | 1079 | | 5851 | 29300748 |
| iAM_Pv461 | *Plasmodium vivax Sal-1* | 461 | 909 | 1078 | | 126793 | 29300748 |
| iAM_Pf480 | *Plasmodium falciparum 3D7* | 480 | 909 | 1083 | | 36329 | 29300748 |
| iLJ478 | *Thermotoga maritima MSB8* | 482 | 570 | 652 | | 243274 | 19762644 |
| iNF517 | *Lactococcus lactis subsp. cremoris MG1363* | 516 | 650 | 754 | | 416870 | 23974365 |
| iSB619 | *Staphylococcus aureus subsp. aureus N315* | 619 | 655 | 743 | | 158879 | 15752426 |
| iJN678 | *Synechocystis sp. PCC 6803* | 622 | 795 | 863 | | 1148 | 22308420 |
| iHN637 | *Clostridium ljungdahlii DSM 13528* | 637 | 698 | 785 | | 748727 | 24274140 |
| iNJ661 | *Mycobacterium tuberculosis H37Rv* | 661 | 826 | 1025 | | 83332 | 17555602 |
| iAF692 | *Methanosarcina barkeri str. Fusaro* | 692 | 628 | 690 | | 269797 | 16738551 |
| iCN718 | *Acinetobacter baumannii AYE* | 709 | 888 | 1015 | | 509173 | 29692801 |
| iJN746 | *Pseudomonas putida KT2440* | 746 | 909 | 1056 | | 160488 | 18793442 |
| iND750 | *Saccharomyces cerevisiae S288c* | 750 | 1059 | 1266 | | 559292 | 15197165 |
| iJB785 | *Synechococcus elongatus PCC 7942* | 785 | 768 | 849 | | 1140 | 27911809 |
| iPC815 | *Yersinia pestis CO92* | 815 | 1552 | 1961 | | 214092 | 21995956 |

| Model ID | Organism | Gene count | Metabolite count | Reaction count | 62 *E.coli* | Taxonomy ID | PubMed ID |
|---|---|---|---|---|---|---|---|
| iSynCJ816 | *Synechocystis sp. PCC 6803* | 816 | 928 | 1044 | | 1148 | 10.1016/j.algal.2017.09.013 |
| iYO844 | *Bacillus subtilis subsp. subtilis str. 168* | 844 | 991 | 1250 | | 224308 | 17573341 |
| iYS854 | *Staphylococcus aureus subsp. aureus USA300_TCH1516* | 866 | 1335 | 1455 | | 451516 | 30625152 |
| iCN900 | *Clostridioides difficile 630* | 900 | 885 | 1229 | | 272563 | |
| iJR904 | *Escherichia coli str. K-12 substr. MG1655* | 904 | 761 | 1075 | | 511145 | 12952533 |
| iMM904 | *Saccharomyces cerevisiae S288c* | 905 | 1226 | 1577 | | 559292 | 19321003 |
| iAF987 | *Geobacter metallireducens GS-15* | 987 | 1109 | 1285 | | 269799 | 24762737 |
| iEK1008 | *Mycobacterium tuberculosis H37Rv* | 1008 | 998 | 1226 | | 83332 | 29499714 |
| iLB1027_lipid | *Phaeodactylum tricornutum CCAP 1055/1* | 1027 | 2172 | 4456 | | 556484 | 27152931 |
| iSDY_1059 | *Shigella dysenteriae Sd197* | 1059 | 1890 | 2540 | X | 300267 | 24277855 |
| iRC1080 | *Chlamydomonas* | 1086 | 1706 | 2191 | | 3052 | 21811229 |
| iSBO_1134 | *Shigella boydii Sb227* | 1134 | 1910 | 2592 | X | 300268 | 24277855 |
| iSbBS512_1146 | *Shigella boydii CDC 3083-94* | 1147 | 1912 | 2592 | X | 344609 | 24277855 |
| iSFxv_1172 | *Shigella flexneri 2002017* | 1169 | 1918 | 2639 | X | 591020 | 24277855 |
| iSFV_1184 | *Shigella flexneri 5 str. 8401* | 1184 | 1917 | 2622 | X | 373384 | 24277855 |
| iS_1188 | *Shigella flexneri 2a str. 2457T* | 1188 | 1914 | 2620 | X | 198215 | 24277855 |
| iSF_1195 | *Shigella flexneri 2a str. 301* | 1195 | 1917 | 2631 | X | 198214 | 24277855 |
| iYL1228 | *Klebsiella pneumoniae subsp. pneumoniae MGH 78578* | 1229 | 1658 | 2262 | | 272620 | 21296962 |
| iSSON_1240 | *Shigella sonnei Ss046* | 1240 | 1938 | 2694 | X | 300269 | 24277855 |
| iAF1260 | *Escherichia coli str. K-12 substr. MG1655* | 1261 | 1668 | 2382 | | 511145 | 17593909 |
| iAF1260b | *Escherichia coli str. K-12 substr. MG1655* | 1261 | 1668 | 2388 | | 511145 | 19840862 |
| iECH74115_1262 | *Escherichia coli O157:H7 str. EC4115* | 1262 | 1918 | 2695 | X | 444450 | 24277855 |
| STM_v1_0 | *Salmonella enterica subsp. enterica serovar Typhimurium str. LT2* | 1271 | 1802 | 2545 | | 99287 | 21244678 |
| iECED1_1282 | *Escherichia coli ED1a* | 1279 | 1929 | 2707 | X | 585397 | 24277855 |
| iECUMN_1333 | *Escherichia coli UMN026* | 1332 | 1935 | 2741 | X | 585056 | 24277855 |

| Model ID | Organism | Gene count | Metabolite count | Reaction count | 62 *E.coli* | Taxonomy ID | PubMed ID |
|---|---|---|---|---|---|---|---|
| iG2583_1286 | *Escherichia coli O55:H7 str. CB9615* | 1283 | 1919 | 2705 | X | 701177 | 24277855 |
| iE2348C_1286 | *Escherichia coli O127:H6 str. E2348/69* | 1284 | 1919 | 2704 | X | 574521 | 24277855 |
| iECSP_1301 | *Escherichia coli O157:H7 str. TW14359* | 1299 | 1920 | 2713 | X | 544404 | 24277855 |
| iECNA114_1301 | *Escherichia coli NA114* | 1301 | 1927 | 2719 | X | 1033813 | 24277855 |
| iECs_1301 | *Escherichia coli O157:H7 str. Sakai* | 1301 | 1923 | 2721 | X | 386585 | 24277855 |
| iLF82_1304 | *Escherichia coli LF82* | 1302 | 1940 | 2727 | X | 591946 | 24277855 |
| iECOK1_1307 | *Escherichia coli IHE3034* | 1304 | 1943 | 2730 | X | 714962 | 24277855 |
| iECS88_1305 | *Escherichia coli S88* | 1305 | 1944 | 2730 | X | 585035 | 24277855 |
| ic_1306 | *Escherichia coli CFT073* | 1307 | 1938 | 2727 | X | 199310 | 24277855 |
| iZ_1308 | *Escherichia coli O157:H7 str. EDL933* | 1308 | 1923 | 2722 | X | 155864 | 24277855 |
| iECP_1309 | *Escherichia coli 536* | 1309 | 1943 | 2740 | X | 362663 | 24277855 |
| iUTI89_1310 | *Escherichia coli UTI89* | 1310 | 1942 | 2726 | X | 364106 | 24277855 |
| iNRG857_1313 | *Escherichia coli O83:H1 str. NRG 857C* | 1311 | 1945 | 2736 | X | 685038 | 24277855 |
| iAPECO1_1312 | *Escherichia coli APEC O1* | 1313 | 1944 | 2736 | X | 405955 | 24277855 |
| iEC042_1314 | *Escherichia coli 042* | 1314 | 1926 | 2715 | X | 216592 | 24277855 |
| iUMN146_1321 | *Escherichia coli UM146* | 1319 | 1944 | 2736 | X | 869729 | 24277855 |
| iECABU_c1320 | *Escherichia coli ABU 83972* | 1320 | 1944 | 2732 | X | 655817 | 24277855 |
| iEcHS_1320 | *Escherichia coli HS* | 1321 | 1965 | 2754 | X | 331112 | 24277855 |
| iECIAI39_1322 | *Escherichia coli IAI39* | 1321 | 1957 | 2722 | X | 585057 | 24277855 |
| iECO103_1326 | *Escherichia coli O103:H2 str. 12009* | 1327 | 1958 | 2759 | X | 585395 | 24277855 |
| iECSF_1327 | *Escherichia coli SE15* | 1327 | 1951 | 2743 | X | 431946 | 24277855 |
| iECDH10B_1368 | *Escherichia coli str. K-12 substr. DH10B* | 1327 | 1947 | 2743 | X | 316385 | 24277855 |
| iBWG_1329 | *Escherichia coli BW2952* | 1328 | 1949 | 2742 | X | 595496 | 24277855 |
| iECO111_1330 | *Escherichia coli O111:H- str. 11128* | 1328 | 1959 | 2761 | X | 585396 | 24277855 |
| iECB_1328 | *Escherichia coli B str. REL606* | 1329 | 1953 | 2749 | X | 413997 | 24277855 |
| iEC55989_1330 | *Escherichia coli 55989* | 1330 | 1953 | 2757 | X | 585055 | 24277855 |

| Model ID | Organism | Gene count | Metabolite count | Reaction count | 62 *E.coli* | Taxonomy ID | PubMed ID |
|---|---|---|---|---|---|---|---|
| iECD_1391 | *Escherichia coli BL21(DE3)* | 1333 | 1945 | 2742 | X | 469008 | 24277855 |
| iETEC_1333 | *Escherichia coli ETEC H10407* | 1333 | 1964 | 2757 | X | 316401 | 24277855 |
| iB21_1397 | *Escherichia coli BL21(DE3)* | 1337 | 1945 | 2742 | X | 469008 | 24277855 |
| iEcE24377_1341 | *Escherichia coli E24377A* | 1341 | 1974 | 2764 | X | 331111 | 24277855 |
| iECIAI1_1343 | *Escherichia coli IAI1* | 1343 | 1970 | 2766 | X | 585034 | 24277855 |
| iEC1344_C | *Escherichia coli C* | 1344 | 1934 | 2726 | X | 498388 | 27667363 |
| iEcSMS35_1347 | *Escherichia coli SMS-3-5* | 1347 | 1949 | 2747 | X | 439855 | 24277855 |
| iECSE_1348 | *Escherichia coli SE11* | 1348 | 1957 | 2769 | X | 409438 | 24277855 |
| iEC1349_Crooks | *Escherichia coli ATCC 8739* | 1349 | 1946 | 2756 | X | 481805 | 27667363 |
| iUMNK88_1353 | *Escherichia coli UMNK88* | 1353 | 1971 | 2778 | X | 696406 | 24277855 |
| iECBD_1354 | *Escherichia coli BL21-Gold(DE3)pLysS AG* | 1354 | 1954 | 2749 | X | 866768 | 24277855 |
| iEKO11_1354 | *Escherichia coli KO11FL* | 1354 | 1974 | 2779 | X | 595495 | 24277855 |
| iECO26_1355 | *Escherichia coli O26:H11 str. 11368* | 1355 | 1965 | 2781 | X | 573235 | 24277855 |
| iEC1356_BI23DE3 | *Escherichia coli BL21(DE3)* | 1356 | 1918 | 2740 | X | 469008 | 27667363 |
| iY75_1357 | *Escherichia coli str. K-12 substr. W3110* | 1358 | 1953 | 2760 | X | 316407 | 24277855 |
| iEcDH1_1363 | *Escherichia coli DH1* | 1363 | 1949 | 2751 | X | 536056 | 24277855 |
| iEC1364_W | *Escherichia coli W* | 1364 | 1927 | 2764 | X | 566546 | 27667363 |
| iJO1366 | *Escherichia coli str. K-12 substr. MG1655* | 1367 | 1805 | 2583 | X | 511145 | 21988831 |
| iEcolC_1368 | *Escherichia coli ATCC 8739* | 1368 | 1971 | 2769 | X | 481805 | 24277855 |
| iEC1368_DH5a | *Escherichia coli DH5[alpha]* | 1368 | 1951 | 2779 | X | 668369 | 27667363 |
| iEC1372_W3110 | *Escherichia coli str. K-12 substr. W3110* | 1372 | 1918 | 2758 | X | 316407 | 27667363 |
| iECW_1372 | *Escherichia coli W* | 1372 | 1975 | 2783 | X | 566546 | 24277855 |
| iWFL_1372 | *Escherichia coli W* | 1372 | 1975 | 2783 | X | 566546 | 24277855 |
| iECDH1ME8569_1439 | *Escherichia coli DH1* | 1439 | 1950 | 2756 | X | 536056 | 24277855 |
| iJN1463 | *Pseudomonas putida KT2440* | 1452 | 2153 | 2927 | | 160488 | |
| iML1515 | *Escherichia coli str. K-12 substr. MG1655* | 1516 | 1877 | 2712 | X | 511145 | 29020004 |
| iYS1720 | *Salmonella pan-reactome* | 1707 | 2436 | 3357 | | | 30218022 |

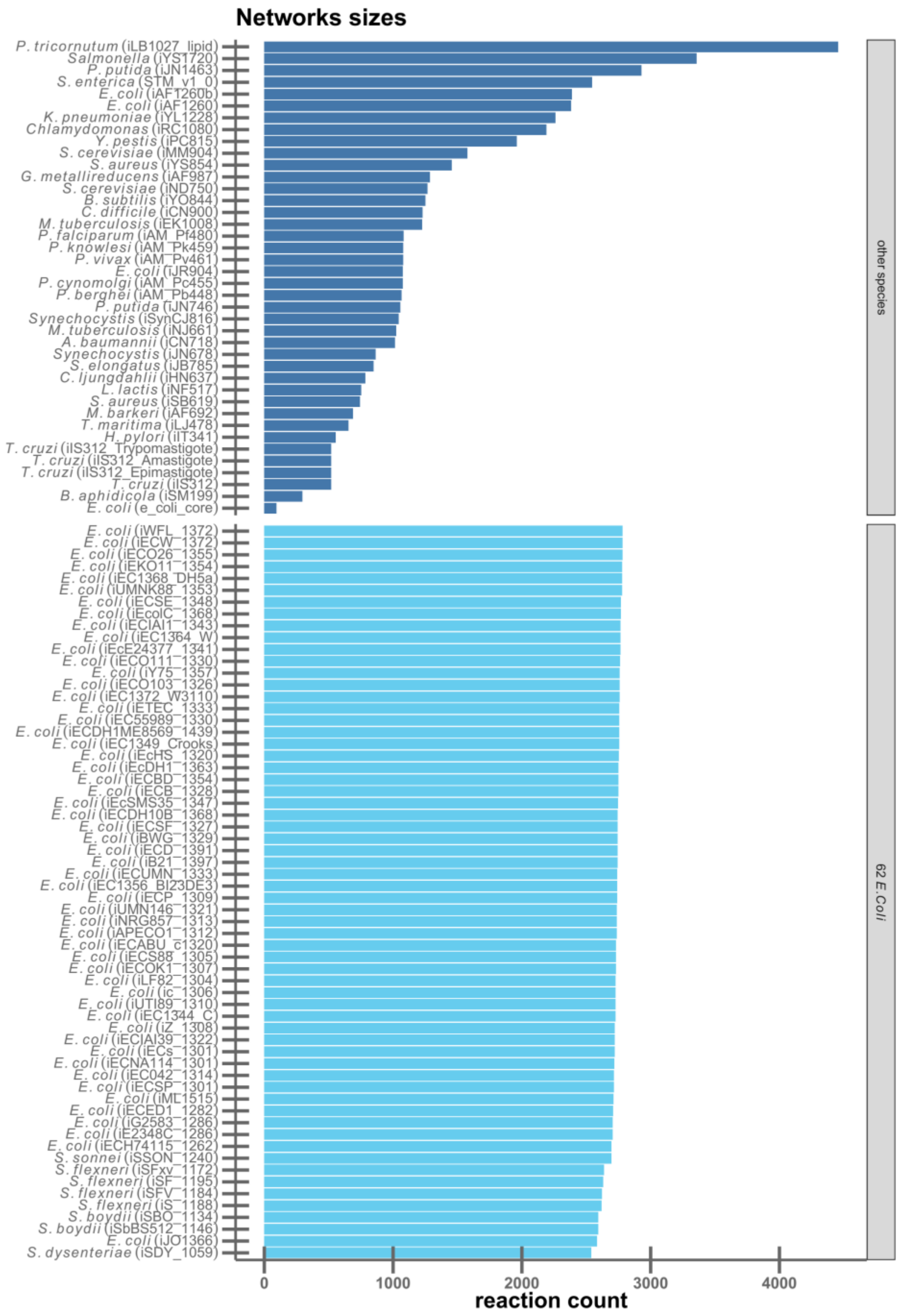

Figure S1. **Networks sizes (number of reactions) of all models combined in the supermodel.** The 62 *E. coli* models shown in an extra group and are depicted in lighter shade. The taxonomy ID refers to the NCBI taxonomy and the PubMed ID refers to the respective publication of the model.

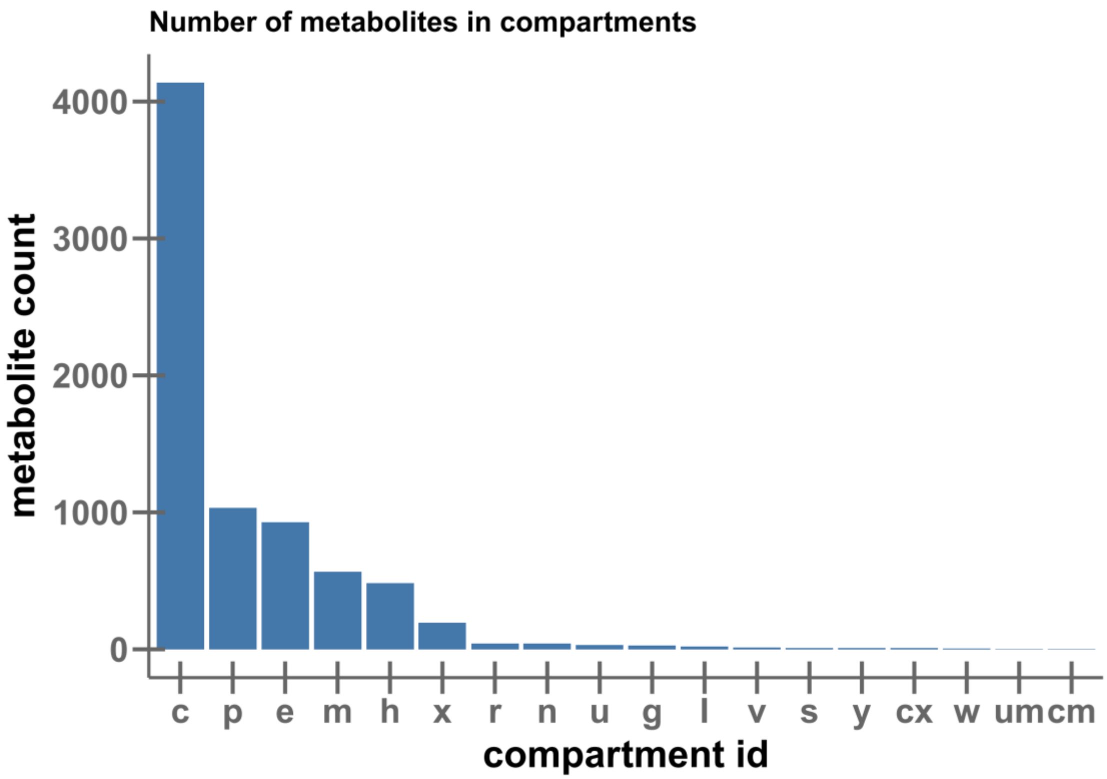

Figure S2. Additional properties of the supermodel. a) Number of metabolites in the compartments. The compartment with the most reactions is the cytosol (c) followed by the extracellular (e) and periplasm (p). The Remaining compartment originate from the eukaryotic models used.

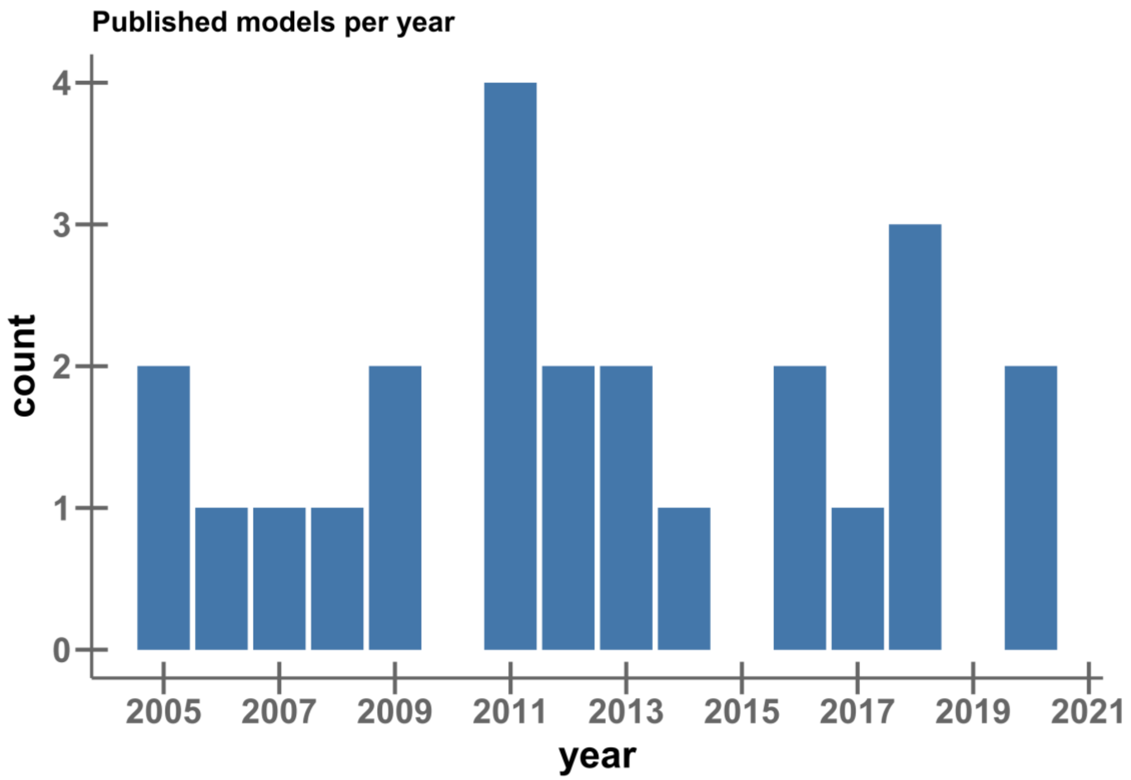

Figure S3**. Bar chart of published models per year.** Here are only the models of unique organisms counted, which were created by about two publications per year.

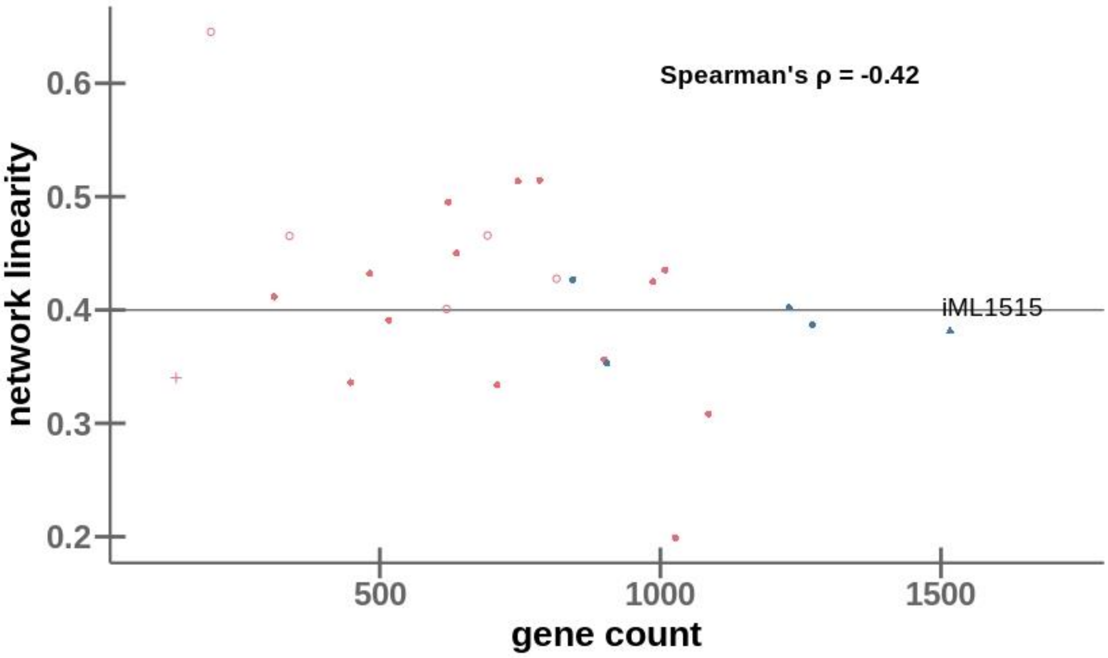

Figure S4. **Small metabolic networks tend to be less branched than large networks.** Network linearity is defined as the fraction of metabolites that participate in only two reactions, i.e., metabolites that are intermediates in unbranched pathways. The colors of circles and points distinguish specialists (red) and generalists (blue). The 62 *E. coli* strains are represented by the iML1515 model (blue triangle) only. Organisms with known auxotrophies are shown as open circles. The highly branched *E. coli* core metabolism is shown with a red plus sign. Spearman correlation between network linearity and gene count: ρ = -0.42, using only iML1515 as representative for the 62 *E. coli*.

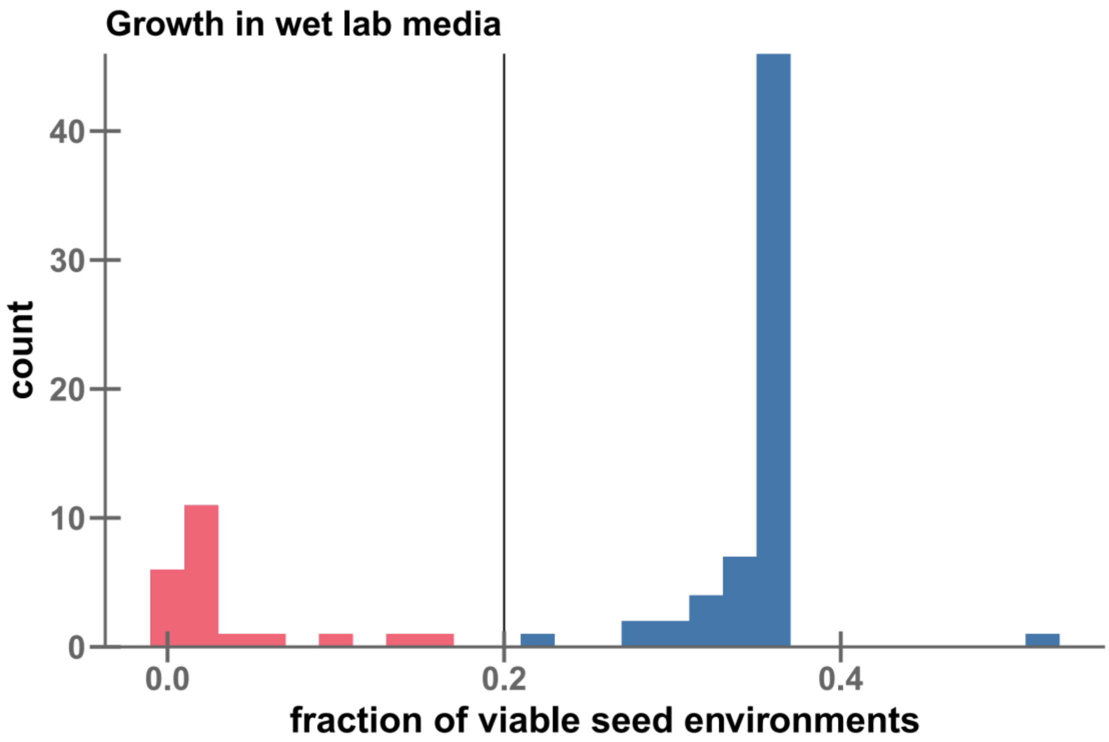

Figure S5. Fractions of viable environments for **submodels in wet lab environments (seed).** The vertical line indicates the threshold to split models into specialists (red) and generalists (blue).

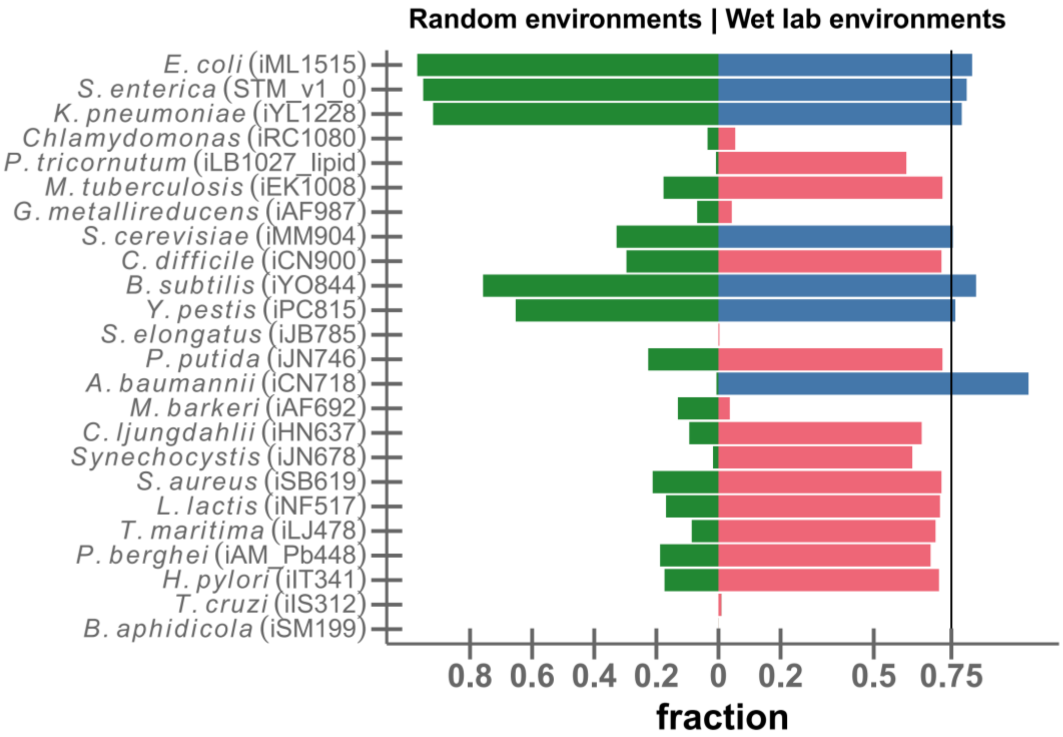

Figure S6. The fraction of viable environments differs widely across submodels, both for random minimal environments (green bars to the left) and for common wet lab environments (blue and red bars to the right), here energy generation as the objective function.

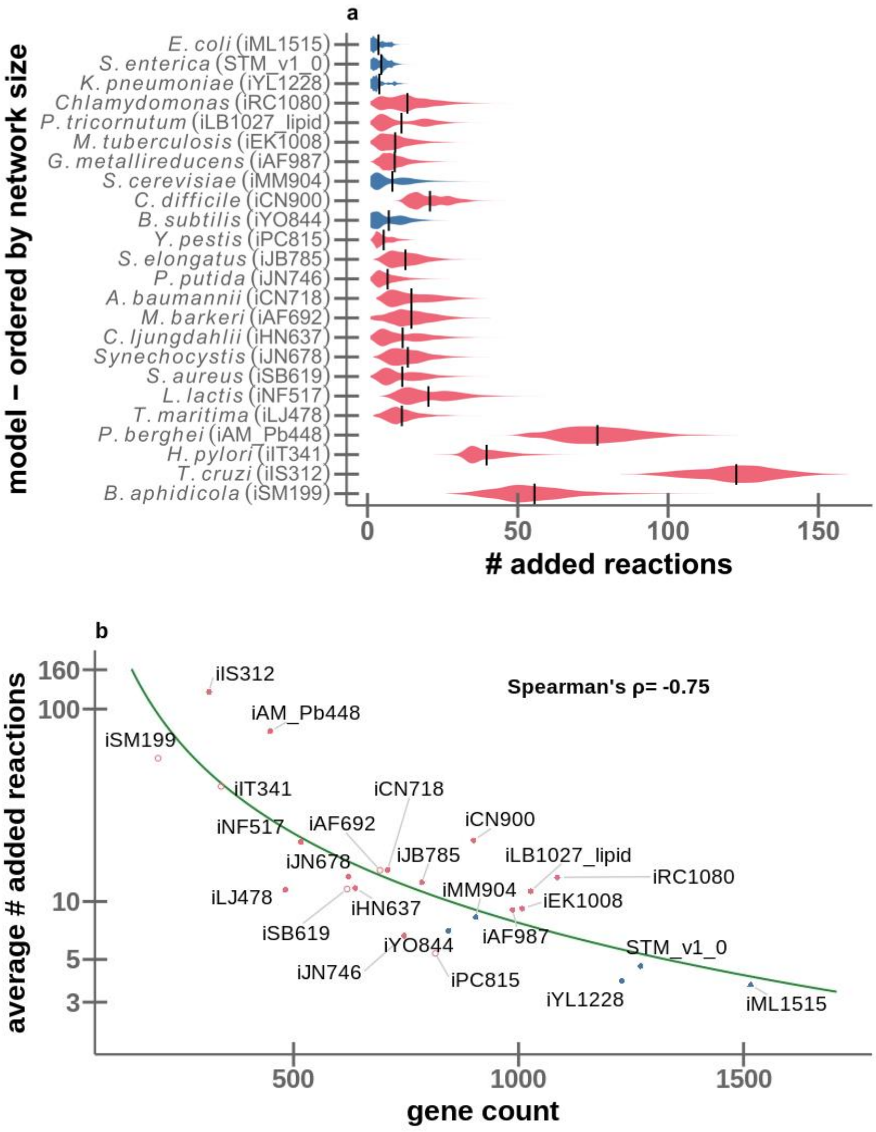

Figure S7. **The number of additional reactions required for adaptation decreases with increasing genome size.** Same as Fig. 3 of the main text, but considering random environments instead of wet lab environments. **(a)** Distributions of added reactions, summarized as violin plots. The height at each point in a "violin" indicates the local density of the distribution for the given model. Models are ordered top-down by decreasing size. **(b)** The average number of added reactions (log scale) plotted against metabolic gene count for each model. The solid line shows the best fitting power law, *added reactions* = $a \times (gene\ count)^b$, with the best-fitting exponent $b$=2.54. In both panels, colors distinguish specialists (red) and generalists (blue). Organisms with known auxotrophies are shown as open circles. The 62 *E. coli* strains are represented by the iML1515 model (blue triangle) only.

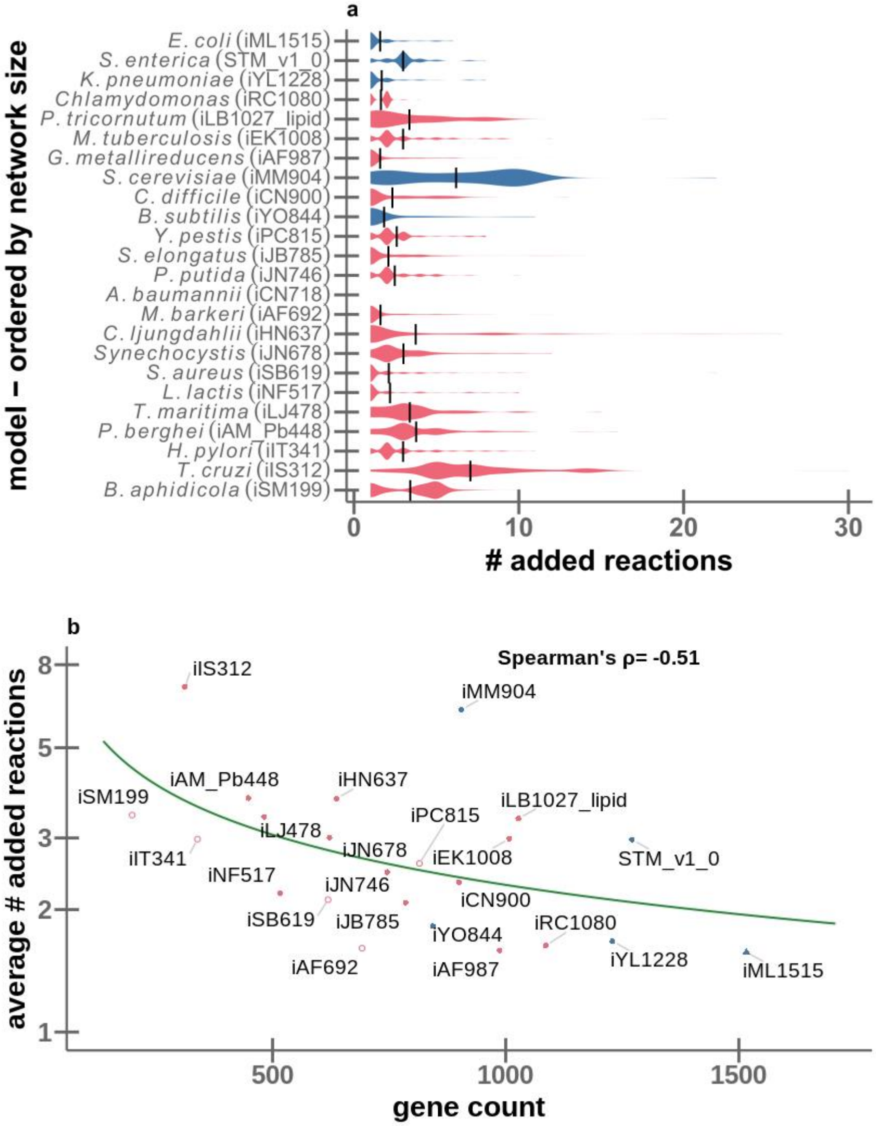

Figure S8. **The number of additional reactions required for adaptation decreases with increasing genome size.** Same as Fig. 3 of the main text, but considering the generation of energy as the objective function (instead of biomass production). **(a)** Distributions of added reactions, summarized as violin plots. The height at each point in a "violin" indicates the local density of the distribution for the given model. Models are ordered top-down by decreasing size. **(b)** The average number of added reactions (log scale) plotted against metabolic gene count for each model. The solid line shows the best fitting power law, *added reactions* = $a \times (gene\ count)^b$, with the best-fitting exponent $b$=1.4. In both panels, colors distinguish specialists (red) and generalists (blue). Organisms with known auxotrophies are shown as open circles. The 62 *E. coli* strains are represented by the iML1515 model (blue triangle) only.

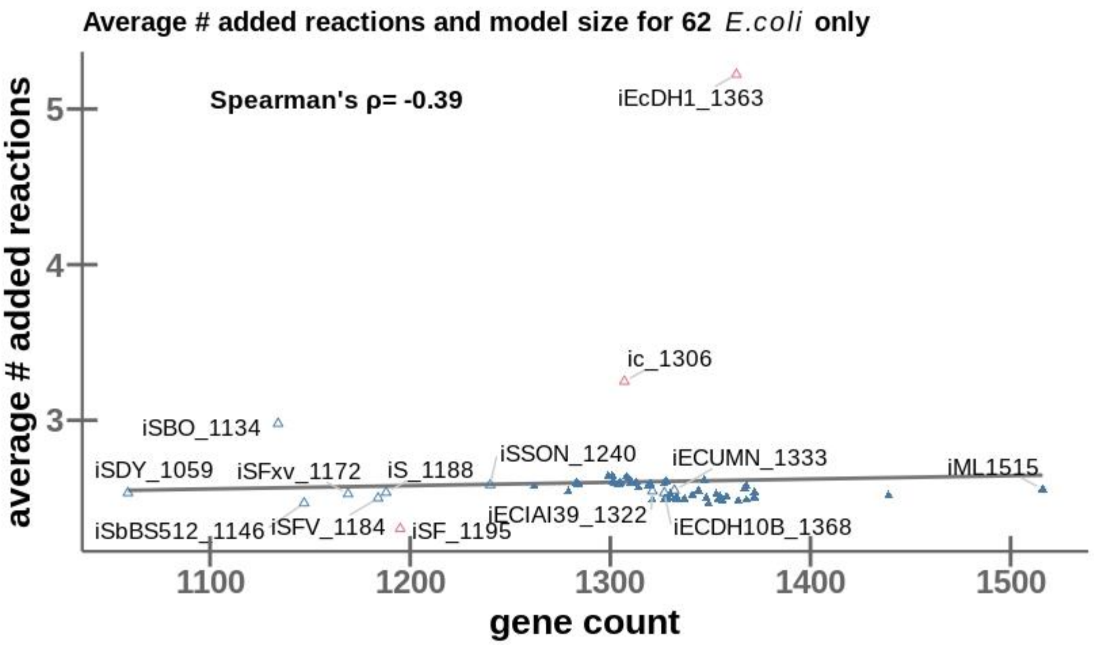

Figure S9. Different E. coli strains show similar adaptabilities, despite variations in genome size. Analogous to Fig. 3b of the main text, but showing all *E. coli* submodels. Colors distinguish specialists (red) and generalists (blue). Organisms with known auxotrophies are shown as open circles.

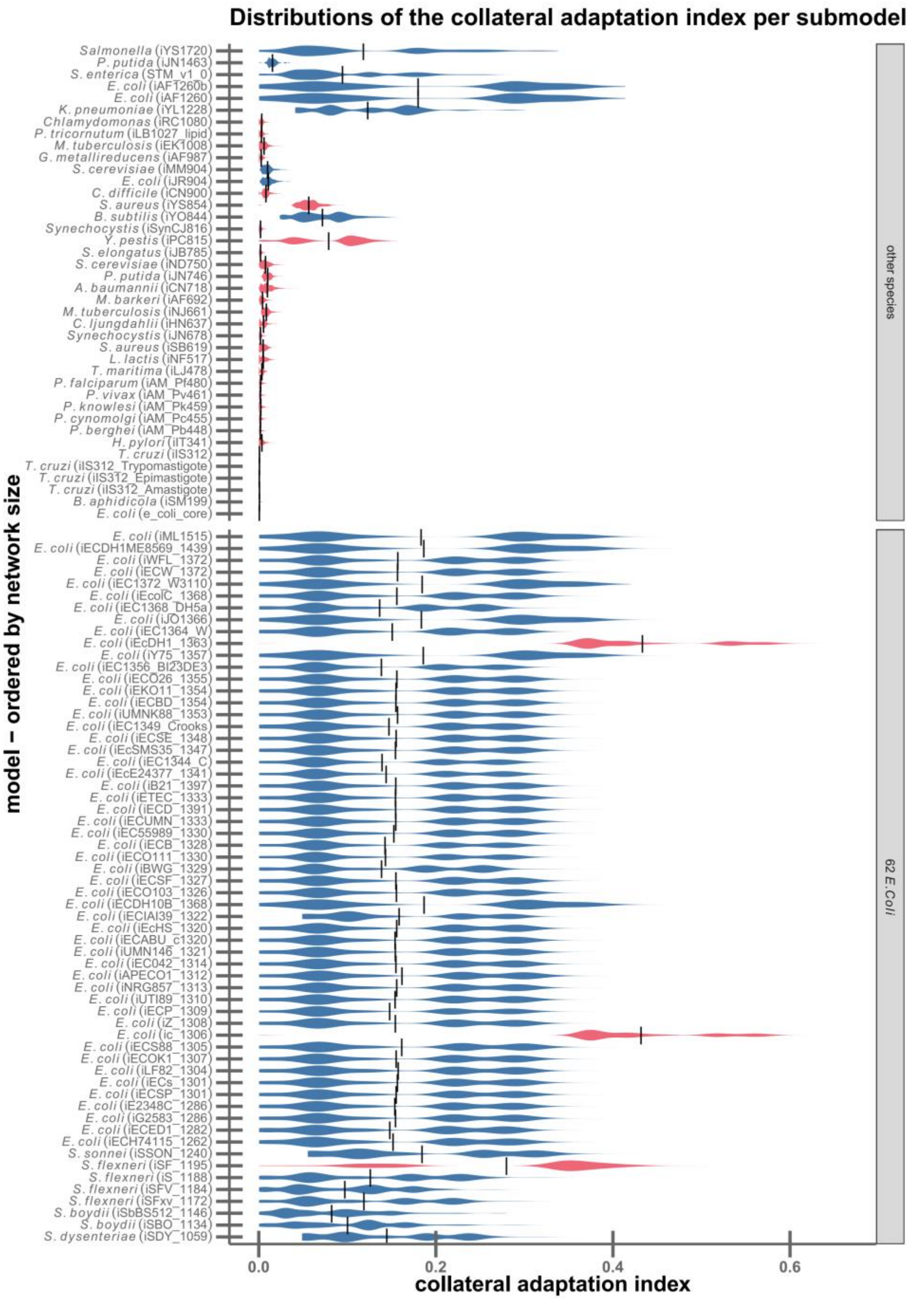

Figure S1**0**. Across species, generalists (blue) show a higher tendency for collateral adaptations than specialists (red), while this trend is reversed within E. coli. For each submodel, we first identified the *n* random environments in which it cannot produce biomass (unviable environments). For each of these environments in turn, we then identified the smallest set of reactions from the supermodel that have to be added to enable biomass production. The collateral adaptation index is then the fraction of the *n*-1 remaining previously unviable environments in which this extended model can grow. Each "violin" summarizes the distribution of the collateral adaptation indices for one submodel. Models in each of the two groups on the y-axis (top: one representative per species; bottom: *E. coli* strains) are sorted by gene count. The mean of each distribution is marked with a vertical line.

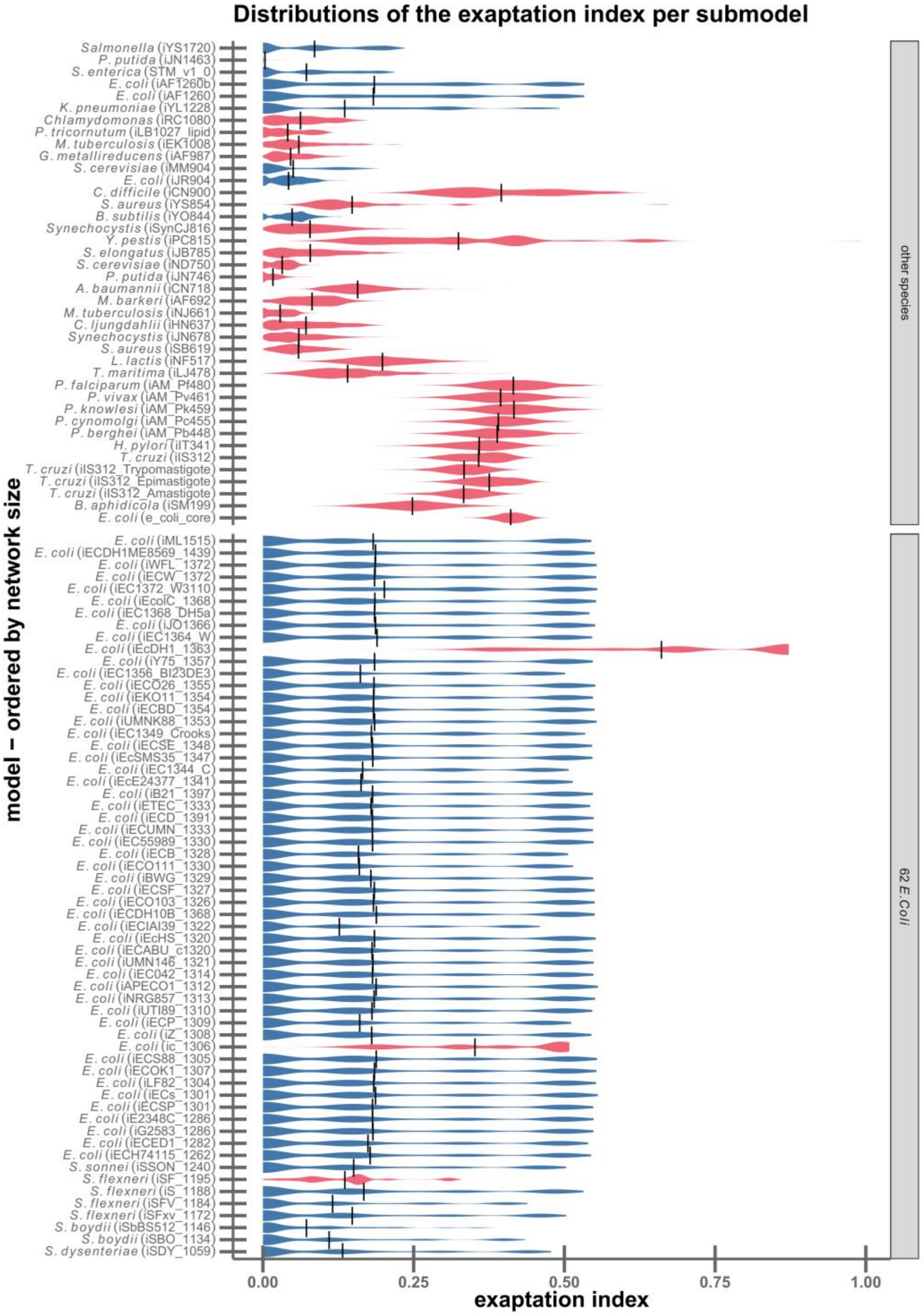

Figure S1**1**. Specialists (red) tend to have higher exaptation indices than generalists (blue). Each "violin" summarizes the distribution of the exaptation index for one submodel. Models in the groups on the y-axis (top: one representative per species; bottom: *E. coli* strains) are sorted by gene count. The mean of each distribution is marked with a vertical line.

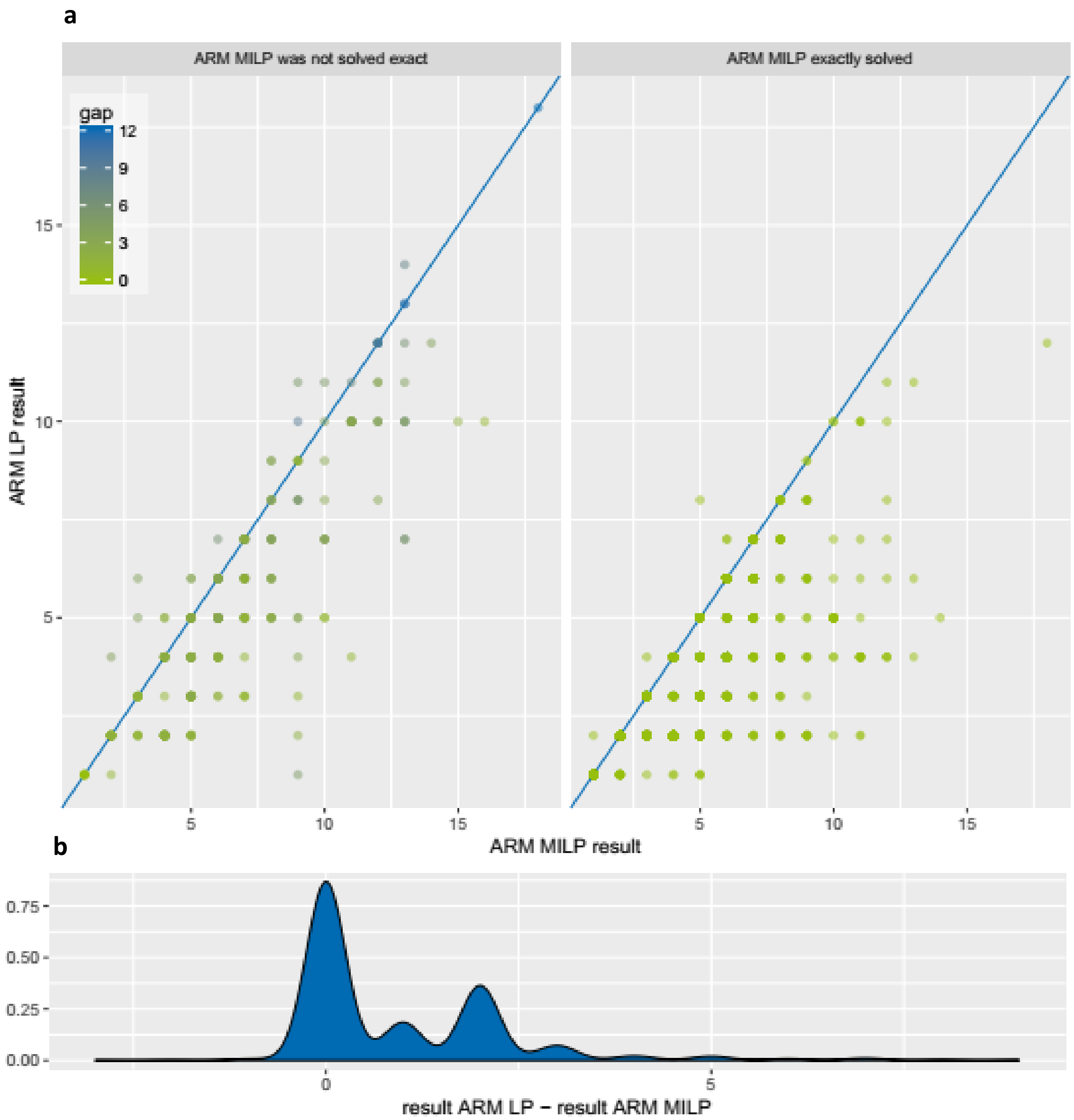

**Figure S12**. Practical application of ARM LP shows equally good performance as ARM MILP. a) Result (objective value) comparison of ARM MILP and ARM LP. Dot color indicates the gap size (smaller is better). In the left panel, ARM MILP solutions are suboptimal due to the limited computation time. Results shown in the right panel could be solved exactly within the time limit. The blue lines indicate equal objective values. b) Distribution of the difference between ARM LP and ARM MILP results

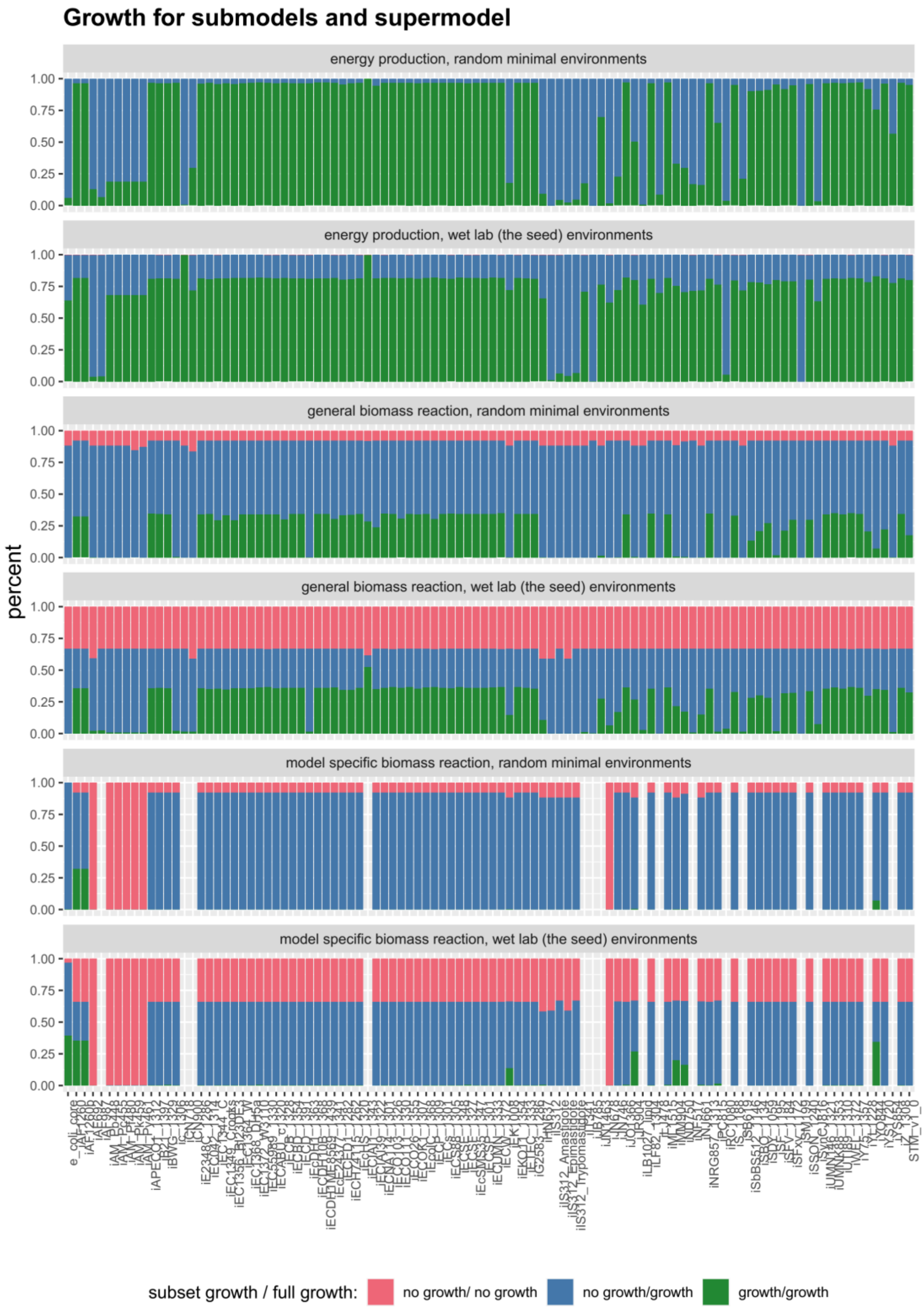

**Figure S13.** Growth for submodels and supermodel in all environment types (random minimal environments, and wet lab (seed) environments) and with three types of biomass objective functions (energy production, general biomass, and organism specific).
